# Resistive force theory and wave dynamics in swimming isolated flagellar apparatus

**DOI:** 10.1101/2020.07.20.211904

**Authors:** S. Goli Pozveh, A. J. Bae, A. Gholami

## Abstract

Cilia-driven motility and fluid transport is ubiquitous in nature and essential for many biological processes, including swimming of eukaryotic unicellular organisms, mucus transport in airway apparatus or fluid flow in brain. The-biflagellated micro-swimmer *Chlamydomonas reinhardtii* is a model organism to study dynamics of flagellar synchronization. Hydrodynamic interactions, intracellular mechanical coupling or cell body rocking are believed to play crucial role in synchronization of flagellar beating in green algae. Here, we use freely swimming intact flagellar apparatus isolated from wall-less strain of *Chlamydomonas* to investigate wave dynamics. Our analysis in phase coordinates show that, when the frequency difference between the flagella is high, neither mechanical coupling via basal body nor hydrodynamics interactions are strong enough to synchronize two flagella, indicating that beating frequency is controlled internally by the cell. We also examined the validity of resistive force theory for a flagellar apparatus swimming freely in the vicinity of a substrate and found a quantitative agreement between experimental data and simulations with drag anisotropy of ratio 2. Finally, using a simplified wave form, we investigated the influence of phase and frequency differences, intrinsic curvature and wave amplitude on the swimming trajectory of flagellar apparatus. Our analysis shows that by controlling phase or frequency differences between two flagella, steering can occur.

## 1. Introduction

Cilia and flagella are hair-like organelles, which protrude from the surface of many eukaryotic cells and play a fundamental role in signal processing [1], sensing [2, 3], propulsion of micro-organisms [4–6] and micro-scale fluid transport [7, 8] at low Reynolds number regime. Cilia and flagella are composed of a microtubule-based structure called the axoneme, and the stresses that lead to the whipping motion are due to dynein molecular motors that are exerting forces between the microtubule doublets by sliding them with respect to each other, converting chemical energy to mechanical work. Constrains at the basal region and along the contour length of axoneme, convert sliding to bending deformations [9–11]. Coordinated beating activity of flagella is crucial for efficient swimming of many ciliated cells in ambient fluid. In response to external stimuli such as light, nutrients, temperature, etc., swimmers transiently change their beating patterns to achieve an efficient directed motion towards the source.

The synchronization mechanism between two or more flagella has been an interesting topic in past and recent years, attracting attention of both physicists and biologists. Synchronized beating patterns of two flagella in single-celled microorganisms such as biflagellate green alga *Chlamydomonas reinhardtii* is necessary for fast directional swimming motion [12]. The two flagella of *C. reinhardtii*, typically beat in synchrony for long time intervals before being interrupted by abrupt large reorientations [12, 13]. It is commonly discussed that interflagellar hydrodynamic interactions between two beating flagella can synchronize their rhythmic patterns [14–20]. However, experiments in Ref. [21] with *C. reinhardtii* measured hydrodynamic forces required for synchronization of two flagella by applying oscillatory external flows and showed that the force is more than one order of magnitude larger than hydrodynamic forces experienced in physiological conditions. Furthermore, it has been shown [22, 23] that when the two flagella desynchronize, the rocking of the cell body brings the beating back to synchrony and the contribution of hydrodynamic coupling is negligible in synchronizing mechanism. However, *C. reinhardtii* cells held fixed with pipette, are also able to synchronize robustly their flagella [13, 24], thus indicating that synchronization is perhaps due to mechanical coupling via internal connecting fibers [25, 26]. Remarkably, synchronization in mutants of *C. reinhardtii* missing the filamentous connections is pronouncedly different from wild types [27]. Other experiments with *C. reinhardtii* [5, 12, 28, 29] also support the crucial role of mechanical coupling through basal bodies [21, 25]. These fibers have a microtubule-based structure showing periodic striation patterns [25]. The periodicity (around 80 nm in *C. reinhardtii*) can change in response to chemical stimuli such as calcium ions, indicating the contractility of the fibers [30].

On the theoretical side, analyses of a three-sphere model in Refs. [23, 31] have demonstrated that for a free swimmer, synchronization can be achieved in the absence of hydrodynamic interactions solely due to local hydrodynamic drag forces which couple oscillatory motion of two flagella via movement of swimmer. Remarkably, in this toy model, in the absence of free translation and rotation of the swimmer, synchronization of the flagellar phases becomes relatively weak, suggesting that to explain synchronization in micro-pipette experiments elastic coupling at the flagellar base is required.

Experiments with the isolated flagellar apparatus from a wall-less mutant of *C. reinhartii* by Hyams & Borisy [32, 33] have shown that both flagella are able to maintain their beating patterns similar to those found in intact cells. Thus, the presence of cell body or cytoplasm is not necessary to synchronize two flagella and is possibly an intrinsic structural property of basal apparatus. In the flagellar apparatus, the two flagella are connected at an angle forming a V-shape at their basal ends with elastic fibers connecting the two basal bodies [25] are plausible candidates to mechanically couple the oscillatory motion of two flagella and synchronize them. For convenience in microscopy, Hyams & Borisy mainly studied on flagellar apparatus anchored to the debris on the substrate and observed that over 70% show synchronous beating patterns, while the rest beat asynchronously. They also observed transient changes between synchrony to asynchrony.

In this work, we studied synchronization dynamics, using phase contrast microscopy, high-speed imaging rates up to 1000 Hz, and image processing to quantify beating patterns of flagellar apparatus isolated from wall-less mutant of *C. reinhardtii*. In contrast to experiments by Hyams & Borisy [32, 33], we examined free-moving flagellar apparatus where these swimmers can easily translate and rotate while the two flagella beat at two different intrinsic frequencies (~16 % of the mean). Unconstrained swimmer movement couples rhythmically beating flagella and this coupling is strongly influenced if swimmer is constrained in movement by e.g. attaching to a substrate or holding in place via a micro-pipette. The question we wish to address in this study may be stated as follows: can coupling via filamentous fibers, swimmer movement or interflagellar hydrodynamic interactions bring two flagella in synchrony, if there is such a high frequency mismatch? Our phase analysis with freely swimming flagellar apparatus demonstrates that these couplings are too weak to cause frequency entrainment. Phase dynamics of oscillator flagella shows that they effectively act as two isolated pendulums beating at their own intrinsic frequencies. They perturb one anothers phases without ever achieving full synchronization. Furthermore, we used our tracked data to check the validity of resistive-force theory (RFT) which neglects long-range hydrodynamic interactions, and found a quantitative agreement between RFT simulations and experimental data with drag anisotropy of ratio 2. Finally, by using a simplified wave form, we performed simulations and analytical approximations to study the swimming motion of flagellar apparatus. This analysis shows that by controlling frequency or phase differences between two flagella, steering of flagellar apparatus can occur.

## 2. Materials and Methods

### 2.1. Isolation of basal apparatus

We used wall-less mutant of *C. reinhardtii* (strain SAG 83.81) to isolate flagellar apparatus, following the protocol of Hymes & Borisy [32]. Briefly, we grew 1.5 liters cultures of cells in TAP (tris-acetate-phosphate) medium at 25 °C in a 14 hours light-10 hours dark illumination cycle to reach cell density of 10^6^ cells/mL. Cells are centrifuged at room temperature for 15 min at 200g and resuspended in 100 mL of HMDEK1 solution at 0 °C (10 mM HEPES, 5 mM MgSO_4_, 1 mM DTT, 0.5 mM EDTA, 25 mM KCL, pH=7), spun down again at 800g for 5 minutes and finally resuspended in 5 mL of HMDEK1. At this step, a small fraction of cells (2-3 %) released their flagellar apparatus while most of the cells released single flagella. A subsequent centrifugation at 800g for 5 minutes, sedimented the cell bodies keeping flagella apparatus and single isolated flagella in the supernatant. For reactivation with ATP, we mixed the suspension with equal volume of HMDEK2 solution (30 mM HEPES, 5 mM MgSO_4_, 1 mM DTT, 0.5 mM EDTA, 25 mM KCL, pH=7) supplemented with 2 mM ATP at pH=7. Note that isolated flagellar apparatus are able to reactivate without de-membranation, since the membrane terminates in the vicinity of basal region of flagella, leaving open ends for free diffusion of ATP [32, 33]. For observation, 10 *μl* solution was infused into 100 μm deep flow chambers, built from cleaned glass and 100 μm thick double-sided tape. The glass surface was blocked using casein solution (from bovine milk, 2 mg/mL) to avoid attachment of basal apparatus to the substrate.

### 2.2. High precision tracking of flagella apparatus

We recorded phase contrast microscopy images of planar swimming basal apparatus at 1000 fps for a duration lasting multiple beating cycles. Phase contrast images are first inverted in intensity and then mean intensity (obtained by averaging over the entire video) is subtracted to increase the signal to noise ratio [34]. A Gaussian filter is applied to smooth the images. In these processed images, the basal body appears as a bright sphere which complicates the tracking procedure of two flagella. Therefore, we removed the basal body by a thresholding step and performed the tracking for each flagellum separately, using gradient vector flow (GVF) technique [35, 36]. In this method, for the first frame, we select a region of interest which contains basal apparatus with two flagella (see Fig. S1). Then, we initialize the snake by drawing a line polygon along the contour of one flagellum in the first frame. This polygon is interpolated at *N* equally spaced points and used as a starting parameter for the snake. The GVF is calculated using the GVF regularization coefficient μ = 0.1 with 20 iterations from the image smoothed by a Gaussian filter. The snake is then deformed according to the GVF where we have adapted the original algorithm by Xu and Prince for open boundary conditions [35, 36]. We repeated this procedure separately for each flagellum which gives us positions of *N* points along the contour length *s* of each flagellum so that *s* = 0 corresponds to the basal end and *s = L* is the distal tip. Here *L* is the contour length of flagellum. The position of each flagellum at *s_i_* is denoted by **r**_1,2_(*s_i_,t*) = (*x*_1,2_(*s_i_, t*), *y*_1,2_(*s_i_, t*)). Indices 1 and 2 refer to first and second flagellum of basal apparatus.

### 2.3. Shape Analysis

The basal apparatus has two flagella, and we performed the mode analysis separately for each flagellum. We described the shape of each flagellum by its unit tangent vector 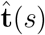 and the unit normal vector 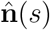 at distance *s* along the contour. Instantaneous deformation of flagella is described by curvature *κ*(*s,t*), which using Frenet-Serret formulas is given by [37]:

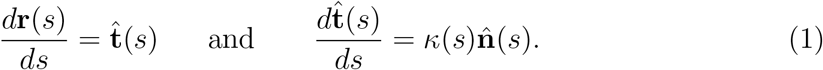

Let us define *θ*(*s*) to be the angle between the tangent vector at distance *s* and the *x*-axis, then *κ*(*s*) = *dθ*(*s*)/*ds*. For shape analysis, we translate and rotate each flagellum such that basal end is at (0, 0) and the orientation of the tangent vector at *s* = 0 is along the 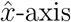 i.e. *θ*(*s* = 0, *t*) = 0. Following Stephens et.al. [37], we performed principal mode analysis by calculating the covariance matrix of angles *θ*(*s, t*) defined as *C*(*s,s′*) = 〈(*θ*(*s,t*) − 〈*θ*〉)(*θ*(*s′,t*) − 〈*θ*〉)〉. We then calculated the eigenvalues *λ_n_* and the corresponding eigenvectors *V_n_*(*s*) of this matrix, showing that superposition of 4 eigenvectors corresponding to four largest eigenvalues can describe the flagella’s shape with high accuracy:

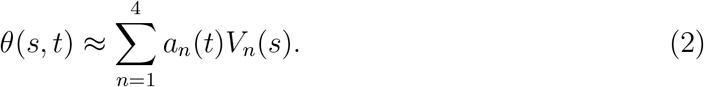

Here the four variables *a*_1_(*t*),.., *a*_4_(*t*) are the amplitudes of motion along different principal components and are given by *a_n_*(*t*) = ∑_*s*_ *V_n_*(*s*)*θ*(*s*).

### 2.4. Resistive Force Theory

As basal apparatus (BA) swims in a fluid, it generates flows of Reynolds number of the order of 10^−3^ or less. Thus, in the absence of inertia, the total force **F**^BA^ and torque **T**^BA^ is zero. The total force **F**^BA^ includes friction forces acting on the basal body **F**^BB^ as well as total hydrodynamics forces on both flagella:

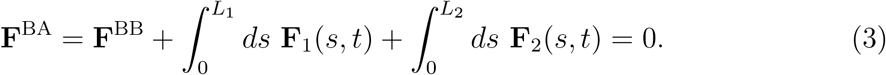

Here *L*_1_ and *L*_2_ are contour lengths of first and second flagella, respectively. Similarly, the total hydrodynamic torque acting on basal apparatus should vanish:

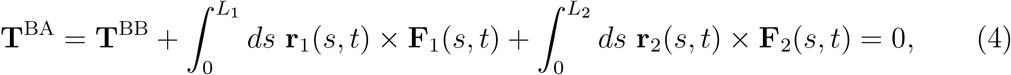

where **T**^BB^ is the torque acting on the basal body. The basal apparatus as a swimmer has two deformable flagella, but at any instant of time it may be considered as a solid body with unknown translational and rotational velocities **U**(*t*) and Ω(*t*) yet to be determined. At small Reynolds number, Newton’s law becomes an instantaneous balance between external and fluid forces and torques exerted on the swimmer:

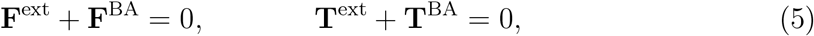

where the force **F**^BA^ and torque **T**^BA^ exerted by the fluid on the basal apparatus are given in Eqs. 3 and 4 and can be separated into propulsive part due to relative deformation of both flagella in body-fixed frame and drag part [38]:

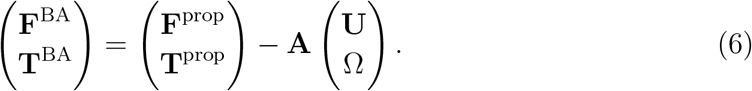

Here *A* is a 6×6 drag coefficients matrix which is symmetric and nonsingular (invertible) and it depends only on geometry of basal apparatus. In our analysis, two flagella and basal body are considered as one long flagellum which is attached to a spherical basal body at its middle. As mentioned above, since in our system, no external forces and torques are acting on basal apparatus, therefore **F**^BA^ and **T**^BA^ must vanish. Furthermore, since basal apparatus swims effectively in 2D, **A** is reduced to a 3×3 matrix and Eq. 6 can be written as:

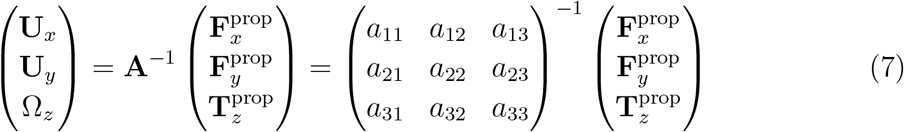

We determine the elements of drag matrix **A** by computing propulsive force and torque exerted by fluid on the swimmer in the lab frame for a translating non-rotating and for a rotating non-translating basal apparatus.

To determine 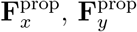 and 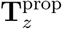 which are propulsive forces and torque due to shape deformations of two flagella in body-fixed frame, we first define a reference frame which is fixed with respect to some arbitrary reference point in basal apparatus, namely the basal body. We set the origin of the body-fixed frame at basal body and define the local tangent vector of first flagellum at contour length *s* = 0 as 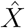 direction, the local normal vector 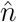 as 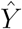 direction, and assume that 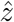 and 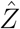 are parallel. We let Φ(*t*) = *θ*(*s* = 0, *t*) to be the angle between 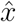 and 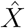 (see Fig. 2A), then in the laboratory frame the velocity of basal body, which was defined as the origin of the body-fixed frame, is given by:

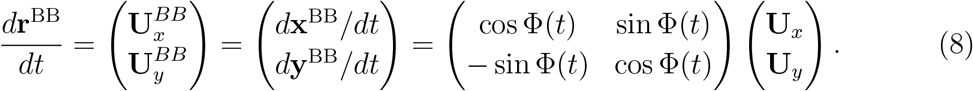

**Figure 2.**
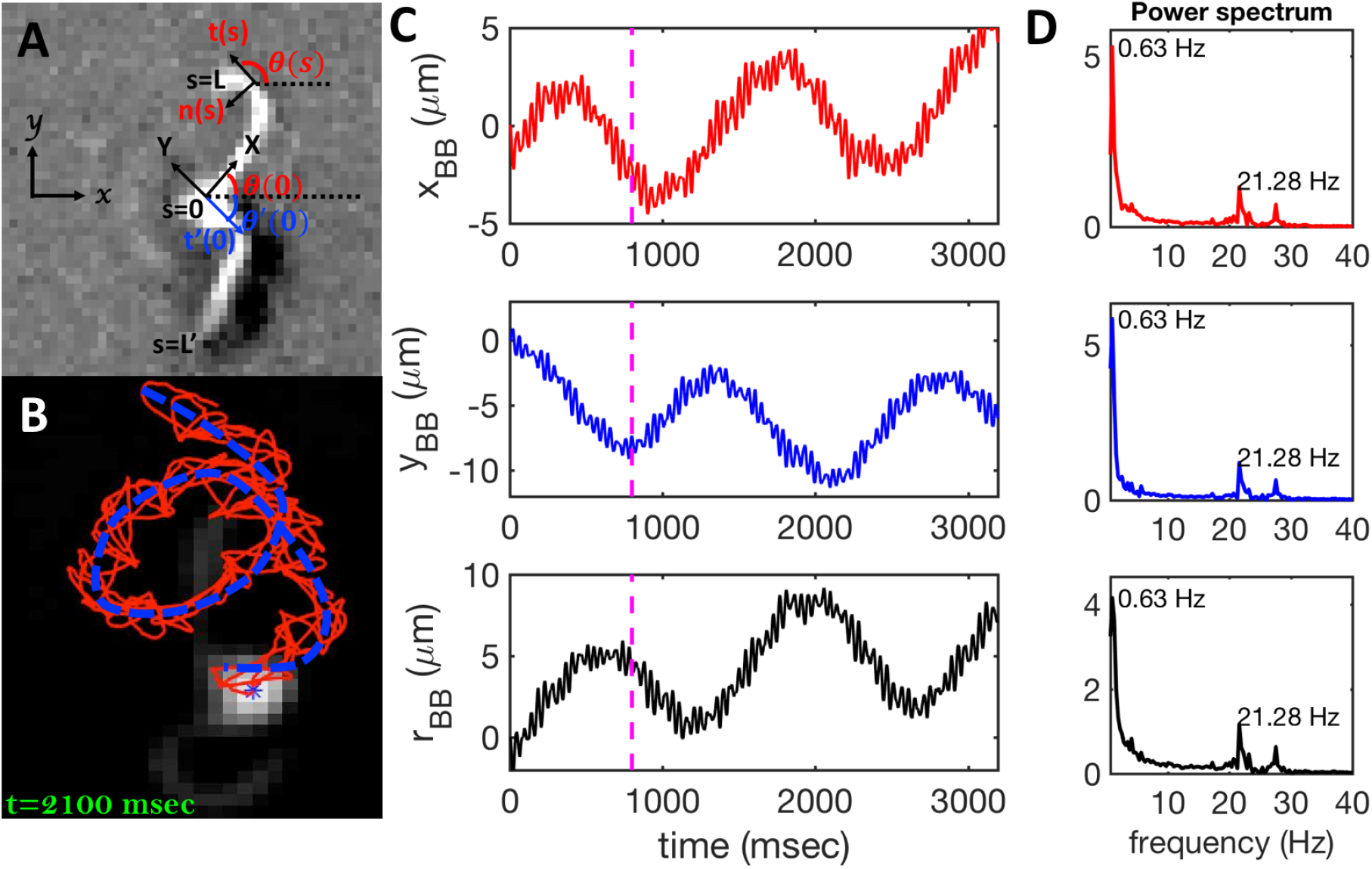
A) Definition of laboratory axes 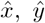 and body-fixed axes 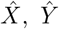. B) Wiggling movement of the basal body on a helical trajectory as it swims with its two flagella in the vicinity of a substrate. B) Displacements of basal body *x_BB_, y_BB_* and *r_BB_* relative to the original position at *t* = 0 showing small-scale oscillations with frequency of 21 Hz (beating frequency of flagella) embedded in large-scale oscillations of frequency around 0.6 Hz. The dashed lines in magenta highlight the first 800 frames that are used for tracking in Fig. 1. D) Power spectrum of *x_BB_, y_BB_* and *r_BB_* displays clear peaks at two frequencies of 21 and 0.6 Hz (see Video 2).

Furthermore, the instantaneous velocity of each flagellum in the lab frame is given by **u = U + Ω × r**(*s, t*) + **u′**, where **u′** is the deformation velocity of flagella in the body-fixed frame, *U* = (**U**_*x*_, **U**_*y*_, 0) and Ω = (0, 0, Ω_*z*_) with Ω_*z*_ = *d*Φ(*t*)/*dt*.

To calculate 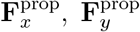 and 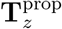 for a given beating pattern of each flagellum in body-fixed frame, we used classical framework of resistive force theory (RFT) which neglects long-range hydrodynamic interactions between different parts of each flagellum as well as inter-flagella interactions. In this theory, each flagellum is divided to small cylindrical segments moving with velocity **u′**(*s, t*) in the body-frame and propulsive force **F**^prop^ is proportional to the local centerline velocity components of each segment in parallel and perpendicular directions:

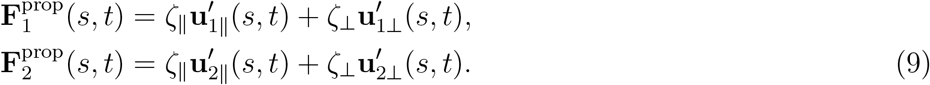

where indices 1 and 2 refer to first and second flagellum in basal apparatus. Here 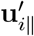 and 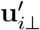 are projections of the local velocity on the directions parallel and perpendicular to each flagellum, i.e. 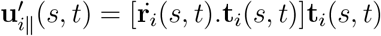 and 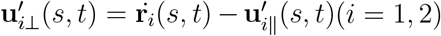 where **t**_1,2_ are the unit tangent vectors along the first and second flagella. The friction coefficients ζ_‖_ and ζ_⊥_ for a thin filament of contour length *L* ~ 10 μm and radius *a ~* 100 nm swimming in surrounding fluid with viscosity μ = 0.96 pN msec/μm^2^ (water at 22°C), are given by ζ_‖_ = 4πμ/(ln(2L/α) + 0.5) ~ 2.1 pN msec/μm^2^ and ζ_⊥_ = 2ζ_‖_. This anisotropy indicates that to obtain the same velocity, one would need to apply a force in the perpendicular direction twice as large as in the parallel direction [39]. Furthermore, in the lab frame, basal ends of both flagella move in synchrony with basal body i.e. 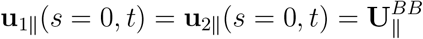 and 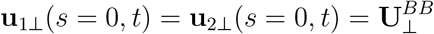.

The effects of basal body connected to the basal ends of both flagella are accounted for by defining **F**^BB^ and **T**^BB^ in Eqs. 3 and 4 to be the drag force and torque acting on the basal body and are given as:

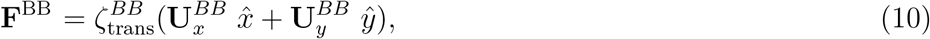

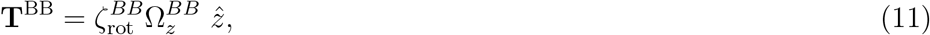

where **U**^*BB*^ and 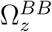 denote translational and rotational speed of basal body in the lab frame. In our analysis, we consider the basal body to be a sphere of dimensionless size *b* = 0.1 which is the ratio between its radius (~ 1 *μm*) and contour length of flagella (~ 10 *μm*). We calculate translational and rotational friction coefficients of basal body as 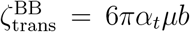 and 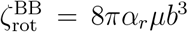, where factors *α_t_* = 16/7 and *α_r_* = 8/7 are corrections due to the fact that basal apparatus swims in the vicinity of a substrate [40].

Here is a short summary of steps in our RFT analysis: first, we translate and rotate basal apparatus such that basal body is at position (0, 0) and the local tangent vector of first flagella at *s* = 0 and time *t* is in the direction defined by tangent vector of first flagella at *t* = 0 and *s* = 0 (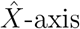 in Fig. 2A). In this way, we lose the orientation information of the basal apparatus at all the time points except for the initial configuration at time t = 0. Note that the isolated flagellar apparatus keeps the V configuration characteristic of the apparatus in situ [25], therefore the angle of V-shape configuration at the basal ends of two flagella which changes over time, is an input from our experimental data used for simulations. Second, we calculate propulsive forces and torque in the body-frame using RFT and Eq. 7, to obtain translational velocities **U**_*x*_, **U**_*y*_ as well as rotational velocity Ω_*z*_ of basal apparatus. Now the rotational matrix can be calculated as:

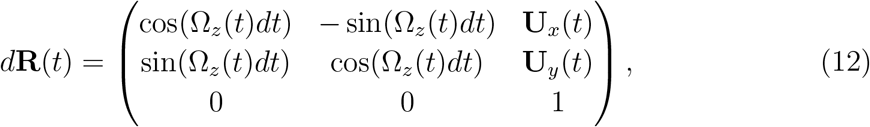

which we use to update rotation matrix as **R**(*t+dt*) = **R**(*t*) ∗ d**R**(*t*), considering **R**(*t* = 0) to be the unity matrix. Having the rotation matrix at time *t*, we obtain the configuration of basal apparatus at time *t* from its shape at body-fixed frame by multiplying the rotation matrix as **r**_lab-frame_(*s, t*) = **R**(*t*) ∗ **r**_body-frame_(*s, t*), which can then be compared with experimental data. Note that **r**_body-frame_(*s, t*) is an input from experimental data presenting the beating patterns in the body-fixed frame.

## 3. Results

### 3.1. Wave form and beating frequencies

Figure 1A illustrates planar swimming motion of an isolated flagellar apparatus immersed in water-like fluid supplemented with 2 mM ATP. If we define the posterior as the basal end of the apparatus, it swims forward as bending waves propagate from basal regions towards the distal tips (Fig. 1B, C). The flagella beat with a power stroke and a recovery stroke comparable to those observed in intact *Chlamydomonas* cells. To characterize the curvature waves and beating frequencies, we tracked each flagellum separately using GVF method (see Fig. 1D and Section. 2.2) and traced the distal ends of both flagella as shown in Fig. 1G. This method gives us *x* and *y* coordinates of N points along each flagellum which is used to calculate the angle between the local tangent vector and 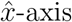, *θ*(*s, t*) (see Fig. 2A).

**Figure 1.**
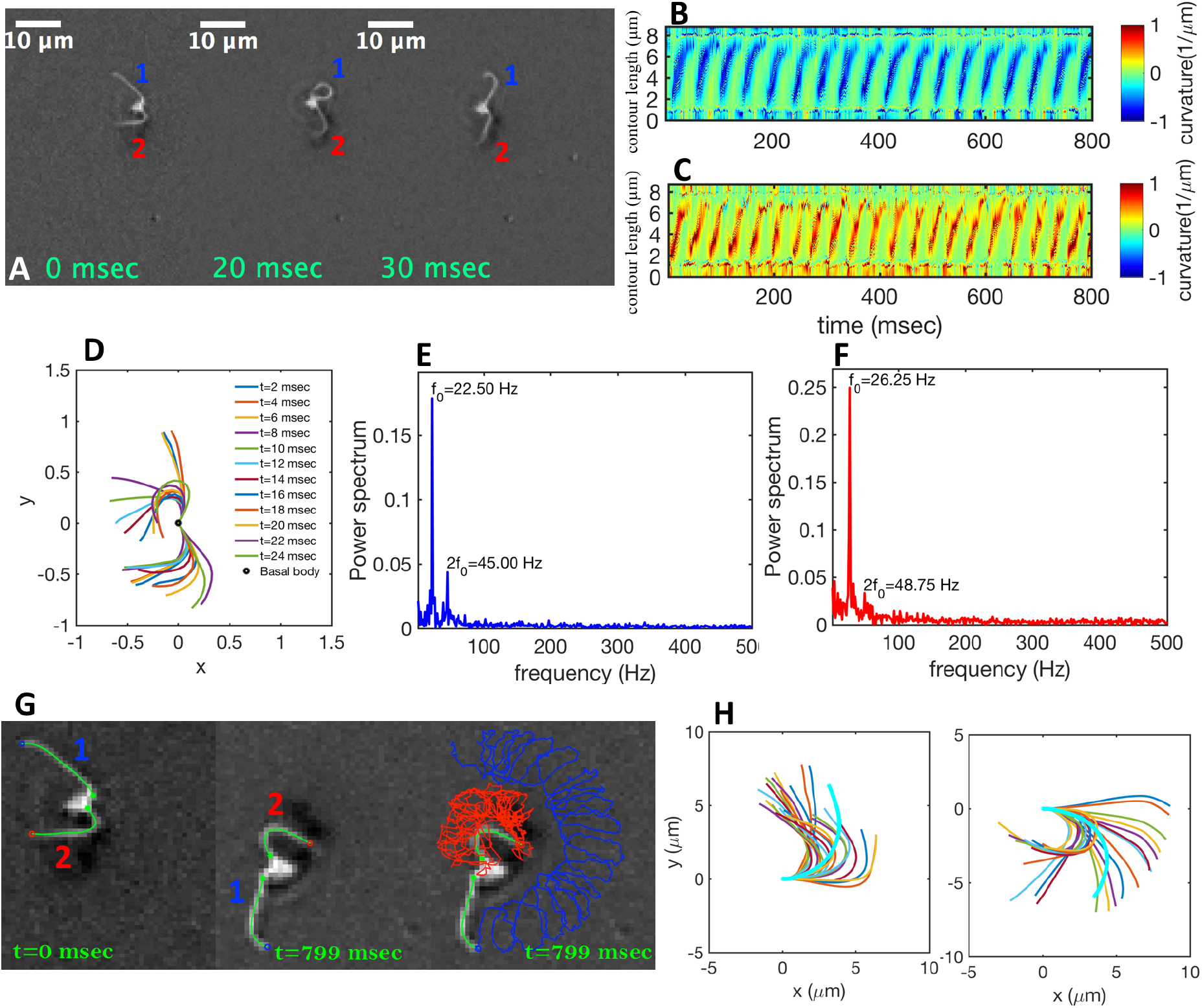
A) Snapshots of swimming isolated flagellar apparatus. B-C) Curvature waves propagating along contour length of both flagella. D) Representative flagellar waveforms, while basal body is translated to be at (0,0). E, F) Power spectral density of both flagella showing that first one beats at the frequency of 22 Hz, while the second one beats faster at frequency of 26 Hz. G) Swimming flagellar apparatus shown at two different time points. Both flagella are tracked using GVF method. Trajectories of distal ends of both flagella as it swims in the time interval of 0 to 799 msec, is shown in the last panel. H) The basal ends of tracked filaments of both flagella are translated to position (0,0) and rotated such that the tangent vector at *s* = 0 is along the x-axis. Semi-circular arcs in cyan color with mean curvature *κ*_0_ ∼ 0.285 *μ*m^−1^ and 0.268 *μ*m^−1^ show the time-averaged shape of flagellum 1 and 2, respectively. This averaged intrinsic curvature makes the waveform asymmetric (see Video 1).

We quantified curvature waves propagating along the contour length of each flagellum using tangent angle *θ*(*s, t*) (Fig. 1B, C). FFT analysis of curvature waves shows dominant peaks at 22.5 Hz and 26.25 Hz for first and second flagellum, respectively (Fig. 1E, F). This is comparable to the typical beating frequencies of flagella in intact wall-less mutants of *Chlamydomonas* cells. We also observed clear peaks at the second harmonics (45 Hz and 48.75 Hz) which temporally breaks the mirror symmetry of beating patterns [41]. Interestingly, oscillatory pattern of each flagellum exhibits a pronounced asymmetry, corresponding to a constant static curvature [42, 43]. We calculated time-averaged shape of each flagellum which results in a semi-circular arc with intrinsic curvature of *κ*_0_/*L* ~ 0.24 rad/*μ*m. Here *L* ~ 10 *μ*m is the contour length of flagellum (see Fig. 1H).

As the flagellar apparatus swims, the basal body follows a helical path as displayed in Fig. 2B. The basal body trajectory shows a wiggle with a frequency 21 Hz embedded in large-scale oscillations with much smaller frequency of 0.63 Hz (Fig. 2C-D). Note that a flagellar apparatus with two flagella beating exactly at the same wave amplitude, frequency and phase, swims in a straight path. However, variations in these parameters generate torque cause the basal apparatus to swim in a circular path (see Sec. 3.3). Even if two flagella beat at the same frequencies, the phase and amplitude of curvature waves are dynamic variables, so the swimmer moving on a curved path does not return precisely to its initial position and follows a helical trajectory.

### 3.2. Mode analysis of flagellar shape and phase dynamics

We performed principal mode analysis to describe time-dependent shape of both flagella in swimming apparatus. This analysis is based on the method introduced by Stephens et.al. [37], which was initially used to characterize wave forms in C. elegance. We used *x* and *y* coordinates of each tracked flagellum, obtained using gradient vector flow technique [35, 36] (see Section 2.2), to calculate *θ*(*s, t*) which is the angle between the local tangent vector of tracked flagellum center line and 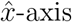 (see Fig. 2A). Next, we computed the covariance matrix *C*(*s, s′*) of fluctuations in angle *θ*(*s, t*) for each flagellum separately. Figure 3A shows the color map of the covariance matrix of the first flagellum which has an effective reduced dimensionality with only small number of nonzero eigenvalues. Remarkably, only four eigenvectors *V_n_*(*s*) (*n* = 1,.., 4) corresponding to the first four largest eigenvalues of *C*(*s,s′*) capture the flagellum’s shape with high accuracy (see Video 3). These four eigenvectors are plotted in Fig. 3B and the first two time-dependent motion amplitudes *a*_1_(*t*) and *a*_2_(*t*) are presented in Fig. 3E.

**Figure 3.**
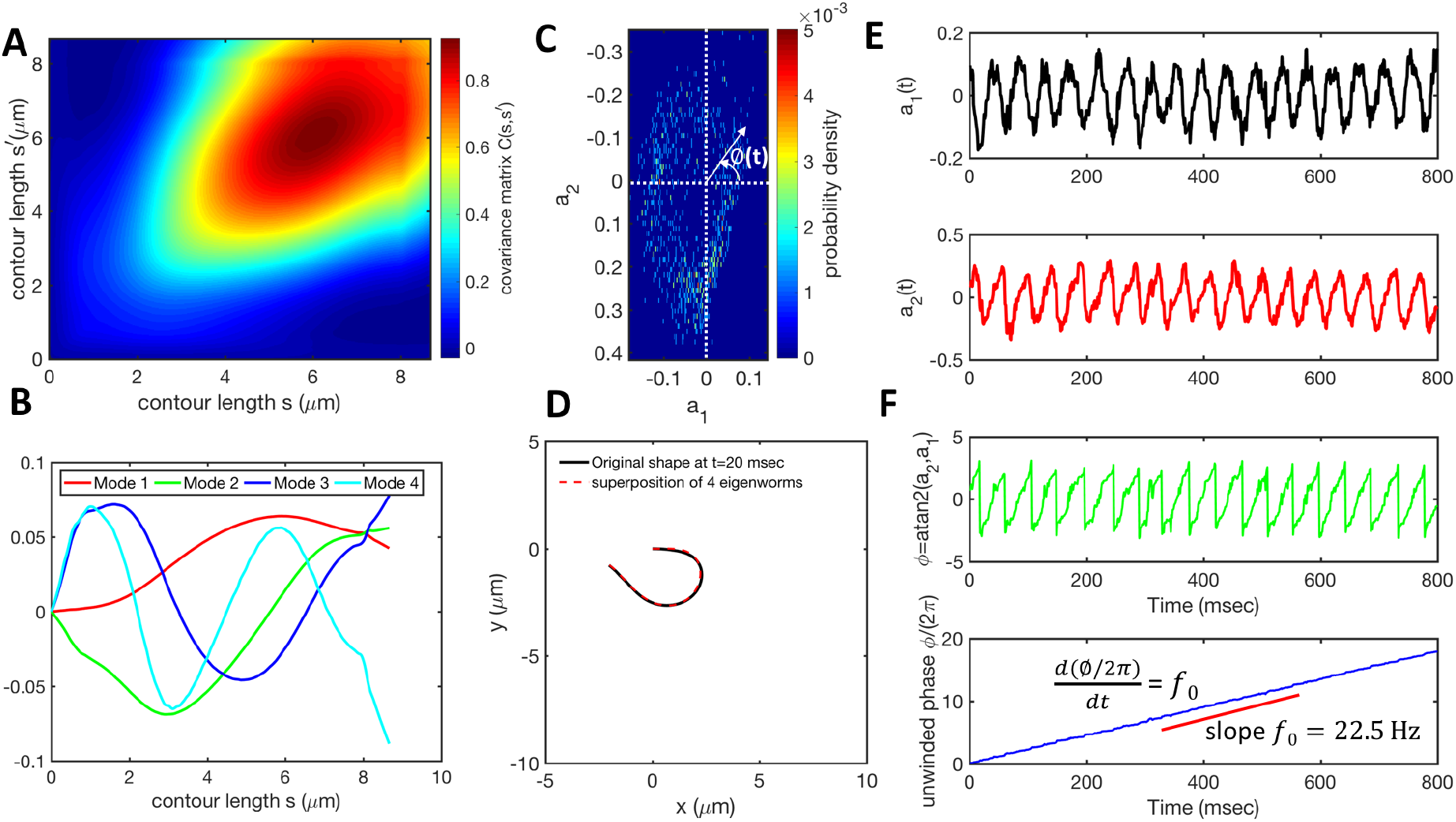
Mode analysis of first flagellum of basal apparatus. Basal end of first flagellum is translated to (0,0) and the tangent angle of the first segment is in 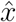 direction. A) The covariance matrix *C*(*s,s′*) of fluctuations in angle *θ*(*s,t*). B) Four eigenvectors corresponding to four largest eigenvalues of matrix *C*(*s, s′*). C) Probability density of the first two shape amplitudes *P*(*a*_1_(*t*), *a*_2_(*t*)). The phase angle is defined as *ϕ*(*t*) = atan2(*a*_2_(*t*), *a*_1_(*t*)). D) Superposition of four eigenmodes presented in part B with coefficients *a*_1_(*t*) to *a*_4_(*t*), can reproduce shape of flagella with high accuracy. E) Time evolution of the first two dominant shape amplitudes *a*_1_(*t*) and *a*_2_(*t*) showing regular oscillations of frequency 22 Hz. F) Dynamics of the phase *ϕ*(*t*) shows linear growth, indicating steady rotation in *a*_1_ − *a*_2_ plane presented in part C. We note that *dϕ/dt* = 2*πf*_0_, where *f*_0_ is the beating frequency.

The Fourier analysis of oscillating motion amplitudes *a*_1_(*t*) and *a*_2_(*t*) gives clear peaks at 22 Hz for first flagellum and 26 Hz for the second one. Figures 1E, F also show the existence of higher harmonics at 45 Hz and 48.75 Hz, respectively [41]. Furthermore, the probability density distribution of *a*_1_(*t*) and *a*_2_(*t*) (Fig. 3C) demonstrates that on average, the principal modes follow a closed trajectory reminiscent of a stable limit cycle, and can be used to define an instantaneous phase as *ϕ*(*t*) = atan2(*a*_2_(*t*), *a*_1_(*t*)). The phase *ϕ* maps variables *a*_1_(*t*) and *a*_2_(*t*) undergoing non-linear oscillations (Fig. 3E), to a new variable which linearly increases over the beating period of each flagellum. By working in phase space, we simply assume that, to leading order, the perturbation due to mechanical coupling via basal body affects only the phase and not the amplitude of the flagella as a non-linear oscillator. Fig. 3F shows the time-dependent phase calculated for first flagellum which is (on average) a monotonically increasing function of time and is equivalent to a uniform rotation in the phase space defined in the *a*_1_ − *a*_2_ plane (Fig. 3C). The time derivative of phase *ϕ*(*t*) is a measure of the oscillation frequency *f*_0_.

The phase dynamics of each flagellum is composed of non-linear fluctuations around a linear deterministic time trend (Fig. 4A). If the two flagella were independent noninteracting oscillators, we would expect the phase difference to grow linearly at a rate proportional to the frequency mismatch Δ*f* (Fig. 4B), and besides some minor fluctuations, this is what we see – i.e. we find no evidence of phase locking and synchronization. Figure 4D illustrates deviation of phase from linear component obtained by subtracting the unwrapped phase of each flagellum plotted in Fig. 4A, from corresponding linear component presented in Fig. 4C. Power spectrum of fluctuations of first flagellum gives clear peaks at 22 Hz and higher harmonics, as shown in Fig. 4E; similarly for the second flagellum.

**Figure 4.**
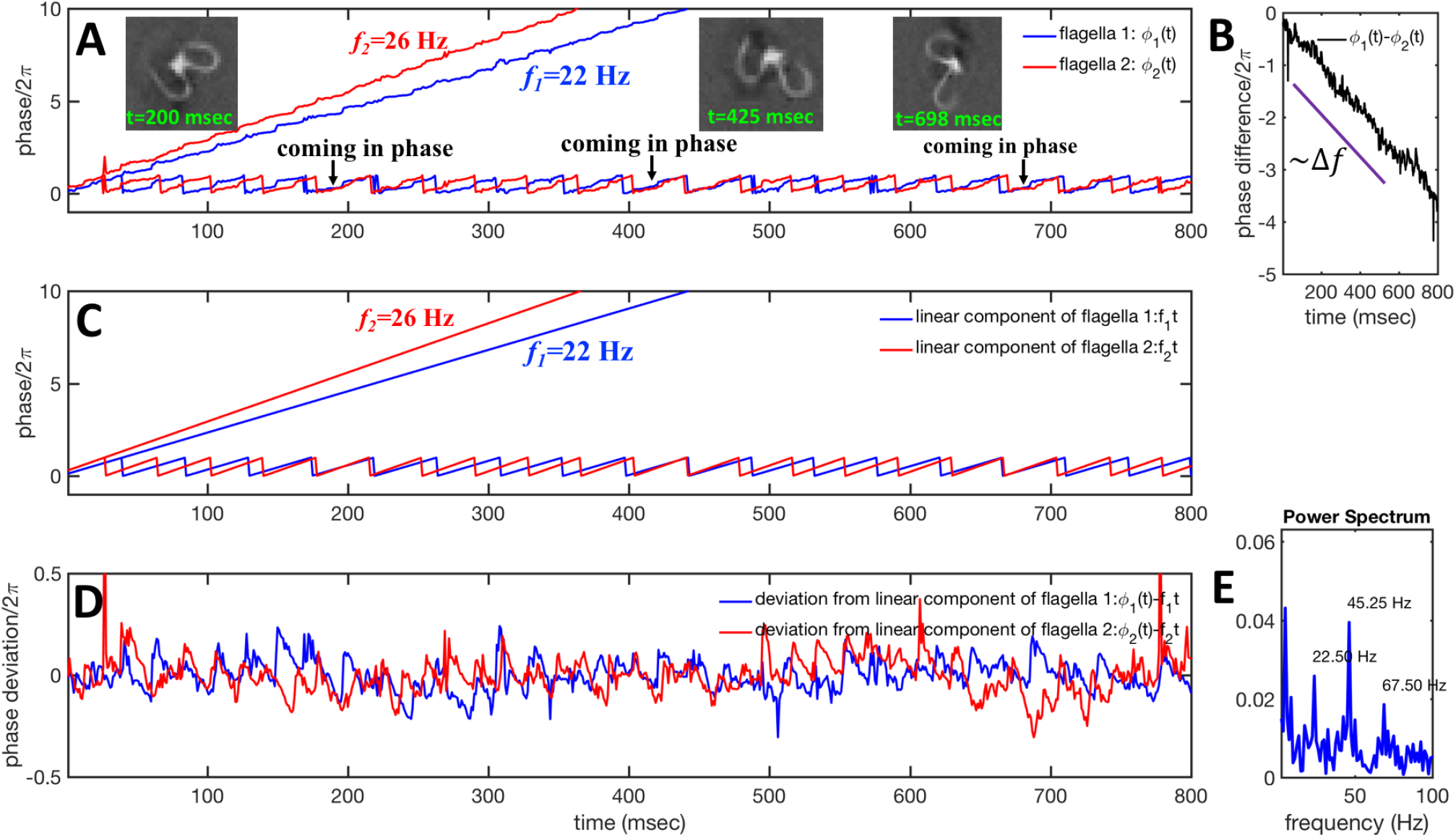
Phase dynamics of two mechanically coupled flagella in swimming basal apparatus. A) For each flagellum, we extract a time-dependent phase *ϕ*(*t*) from motion amplitudes *a*_1_(*t*) and *a*_2_(*t*) defined as *ϕ*(*t*) = atan2(*a*_1_(*t*),*a*_2_(*t*)). Unwrapped phase for each flagellum shows fluctuations around a linear trend. B) For a frequency mismatch of 4 Hz (16% of the mean), the two flagella display phase differences that varies monotonously with time. C) Linear phase component of each flagellum that scales with frequency. D) Deviations from linear components where unwrapped phase of two flagella in part A is subtracted from corresponding unwrapped linear components presented in part C. E) Power spectrum of non-linear fluctuations of first flagellum shows clear peaks at beating frequency of 22.5 Hz and at higher harmonics.

Obviously, mechanical coupling fail to entrain two flagella and on average, they behave as two isolated oscillators with phase evolving at constant rates *ω*_1_ = 2*π f*_1_ and *ω*_2_ = 2*π f*_2_ in time. As two flagella beat with 4 Hz frequency difference, over time they find similar phase values (black arrows in Fig. 4A), corresponding to the time points that two flagella for a short time beat in synchrony. Frequency difference quickly drives the system out of synchronous state to asynchronous phase, which is the dominant state during swimming period of flagellar apparatus (see Video 1).

### 3.3. Quantitative agreement with resistive force theory

We used the experimental beating patterns of the flagellar apparatus to examine the validity of resistive force theory in our *in vitro* model system. At each time point t, the configuration of flagellar apparatus in the lab frame was calculated from instantaneous translational and rotational velocities of flagellar apparatus in the body-fixed frame using the rotation matrix presented in Eq. 12. The control parameter was the drag anisotropy ζ_⊥_/ζ_‖_ which is the ratio between drag coefficients in perpendicular and tangential directions. Given experimental wave forms, we compared RFT simulations with instantaneous translational and rotational velocities of flagellar apparatus in the body-fixed frame (see Figs. 5 and 6). We found a quantitative agreement for drag anisotropy of ζ_⊥_/ζ_‖_ = 2. Note that in the body-fixed frame, the motion of basal body is described by time-dependent translational velocities **U**_*x*_, **U**_*y*_ and rotational velocity Ω_z_ which is a measure of rotation of the basal apparatus. In the lab frame, the translational velocities of basal body are given by 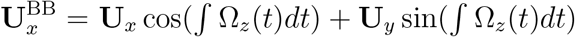 and 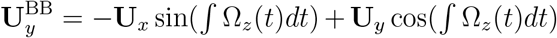. Figure 6 shows *U_x_, U_y_* and Ω_*z*_ obtained from RFT analysis and a comparison with direct experimental measurements in the body-fixed frame, computed by first differentiating experimental *x*_BB_, *y*_BB_ and *θ*(*s* = 0) with respect to time and then transforming to the swimmer-fixed frame. Remarkably, the corresponding power spectra of *U_x_, U_y_* and Ω_*z*_ show dominant peaks at 22.5 Hz indicating that beating frequency of first flagellum is reflected in translational and rotational velocities of basal body.

**Figure 5.**
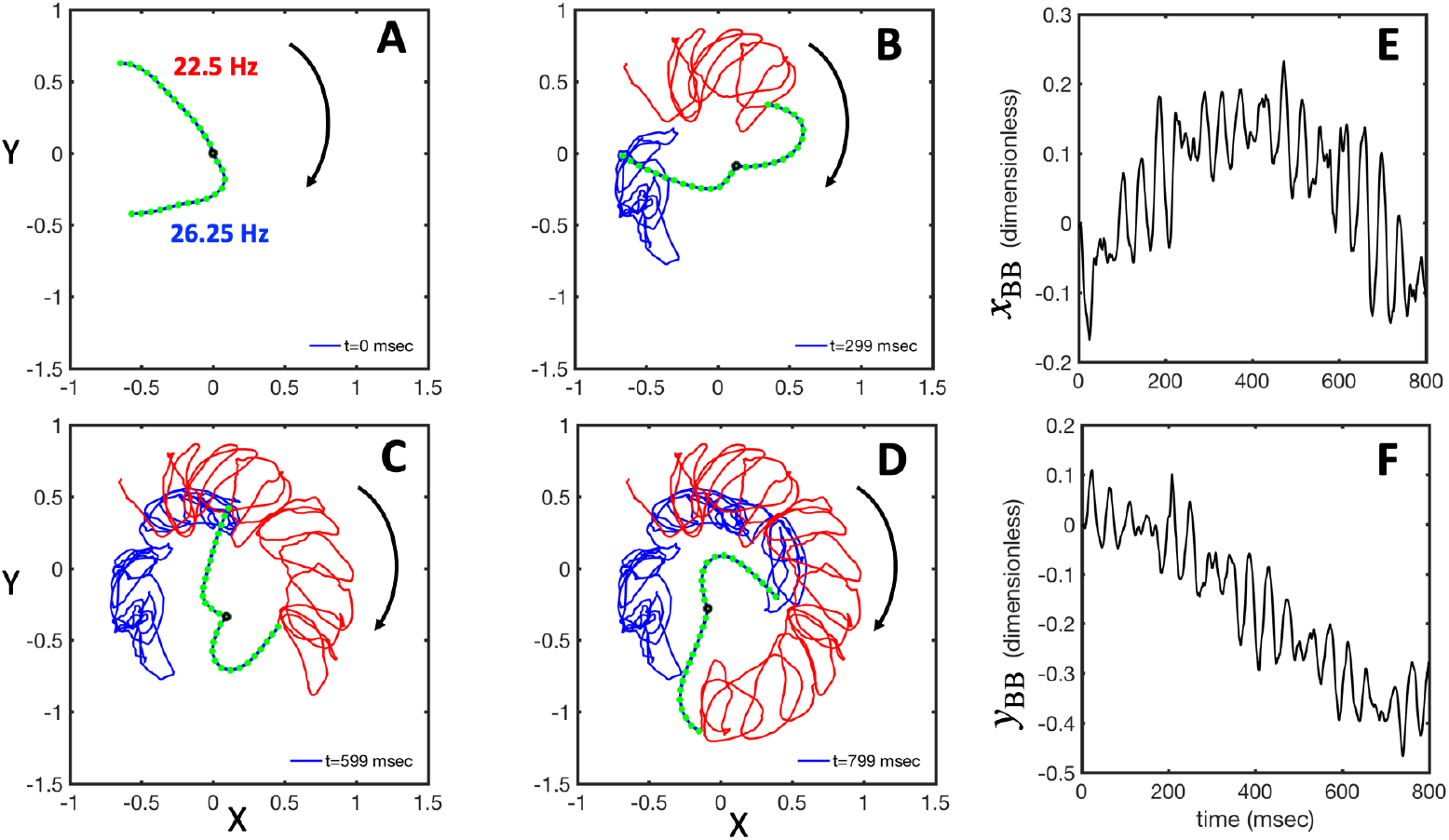
RFT simulations using experimental beating patterns. A) Initial configuration of flagellar apparatus at *t* = 0 extracted from experimental data (compare with Fig. 1G at *t* = 0). B-D) Swimming trajectory of flagellar apparatus obtained by RFT simulations. E-F) Dimensionless positions *x*_BB_ and *y*_BB_ of basal body obtained from RFT, display oscillations reflecting the beating frequency of flagella. Lengths are non-dimensionalized to the contour length of flagella. (see Video 4).

**Figure 6.**
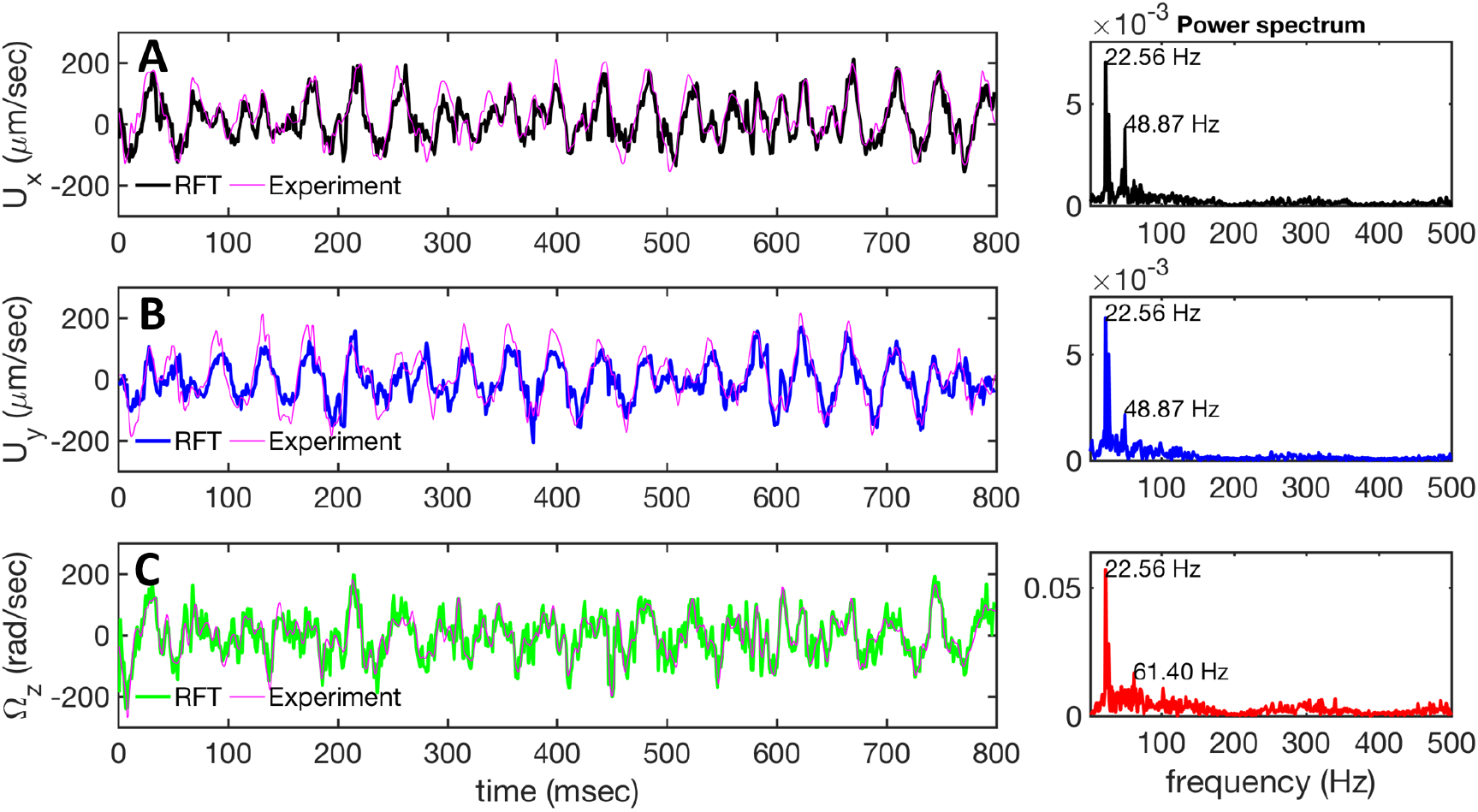
Comparison between RFT analysis and experiments. A-C) Instantaneous translational and rotational velocities of basal apparatus **U**_*x*_, **U**_*y*_ and Ω_*z*_ (*t*) in body-fixed frame, obtained from RFT simulations using experimental beating patterns. Direct experimental results are shown in magenta. Power spectra of translational and angular velocities show clear peaks around 22.5 Hz, which is the beating frequency of first flagellum.

#### Simulations with simplified wave forms

To investigate the effect of wave amplitude, frequency and phase difference of two flagella on the swimming trajectory of the basal apparatus, we performed simulations with simplified form of curvature waves which is a superposition of a dynamic and a static mode [43]:

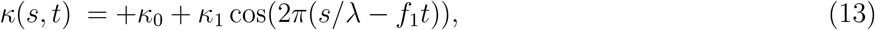

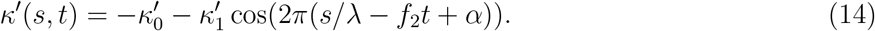

Here λ is the wavelength of curvature waves typically on the order of the flagellar length *L*, 2*πα* is the phase difference between two flagellar beats, and constant intrinsic curvature 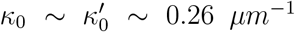 are determined from experimental data by characterizing time-averaged shape of each flagellum, as illustrated in Fig. 1H. Furthermore, *κ*_1_ and 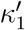 are the amplitudes of the dynamic modes which are estimated from experimental wave patterns to be around 0.45 *μm*^−1^. The negative signs in terms f1t and f2t generate curvature waves that propagate from basal regions (*s* = 0) towards the distal tips (*s = L*), as observed experimentally. Two flagella are positioned exactly symmetrically with respect to the axis of symmetry of the basal apparatus and in our simulations, we assume V-junction angle to be fixed at 180^°^. To perform simulations, we compute drag force density felt by each flagellum in the framework of resistive-force theory (see Eq. 9). The basal body is assumed to have a dimensionless radius 0.1.

In general, the mean rotational velocity 〈Ω_*z*_〉 of flagellar apparatus depends on the amplitudes of static and dynamic modes as well as the frequency and phase difference between the two flagella. For a single flagellum beating at frequency *f*_0_, in the small-curvature approximation assuming *κ*_0_*L*/(2*π*) and *κ*_1_*L*/(2*π*) to be small, (Ω_*z*_) depends linearly on *κ*_0_ but is proportional to the square of *κ*_1_ (see Appendix I and Refs. [41, 44]):

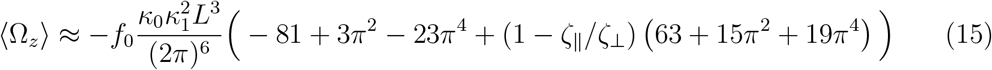

We emphasize that 〈Ω_*z*_〉 is non-zero even if the drag anisotropy is zero (ζ_‖_ = ζ_⊥_). We also note that mean rotational velocity 〈Ω_*z*_〉 scales as *κ*_1_ and therefore is one order of magnitude smaller than instantaneous rotational speed Ω_*z*_(*t*) which scales as *κ*_1_ [44–46].

In the case of flagellar apparatus with two flagella, ignoring hydrodynamic drag force of the basal body for simplicity, rotational velocity has contribution from both flagella (see Appendix I):

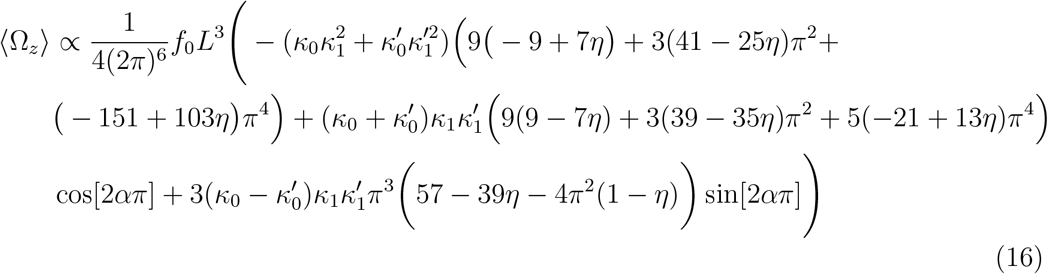

where *η* = 1 − ζ_‖_/ζ_⊥_ and we have assumed the same beating frequency *f*_0_ for both flagella. In the limit of 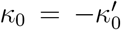 and 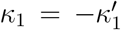, where the two flagella waveforms are mirror images having same parameters with only a phase difference between them, Eq. 16 simplifies to:

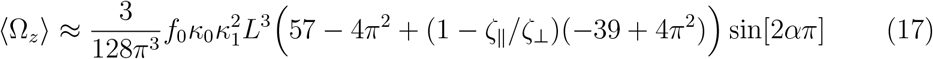

and in the limit of α = 0, Eq. 16 reduces to:

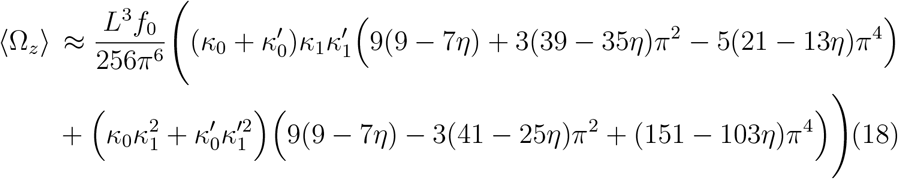

We comment on some properties of Eqs. 16–18: flagellar apparatus with two flagella beating exactly at the same frequency and phase, with mirror-symmetric waveforms (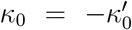 and 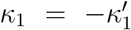), will swim in a straight path with its translational velocity Uy oscillating at frequency of flagellar beat (Fig. 7A, D). However, only a phase difference between two flagella is enough to change the swimming trajectory to a circular path (Fig. 7B-C, E-F). It is noteworthy that the mean rotational velocity 〈Ω_*z*_〉 scales with sin(2*πα*) and thereby, by increasing the phase difference in the range 0 ≤ 2*πα* ≤ *π*/2, swimmer rotates on average faster as illustrated in Fig. 7C. At phase difference between *π*/2 ≤ 2*πα* ≤ *π*, 〈Ω_*z*_〉 starts to decrease and vanishes at *π* (*α* = 1/2) (see Eq. 17). Furthermore, a frequency difference between two flagella also generates a circular swimming path. This is shown in Fig. 8A, where two flagella beat at 50 Hz (red trajectory) and 56 Hz (blue trajectory), while all the other parameters are kept the same. Since in our analytical approximation, to calculate the time-averaged Ω_*z*_, we consider only one single frequency f0 (see Eq. S.14), the influence of having two different beating frequencies is not reflected in our expression in Eq. 16. Another point to mention is that the amplitude of dynamic mode enters as 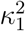 and 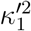 in the Eq. 18 and by introducing a small difference in values of *κ*_1_ and 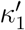, we can switch the swimming direction. Figure 8B shows an example of a flagellar apparatus where the flagellum with red trace has a smaller frequency but larger amplitude of the dynamics mode *κ*_1_, while the other flagellum with blue trace has larger frequency but smaller 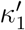. Thus, the flagellum with larger *κ*_1_ wins and sets the sign of (Ω_*z*_). Remarkably, if the beating frequency, amplitude of dynamic mode and phase are equal for both flagella (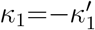 and *α* = 0), then Ω_*z*_ is proportional to 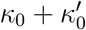:

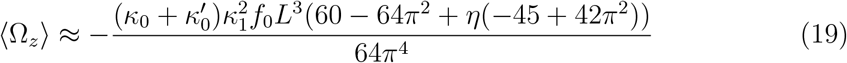

indicating that flagellum with larger intrinsic curvature sets swimming direction on the circular path. Further, Eqs. 16 and 19 show that for an isotropic drag coefficients ζ_‖_ = ζ_⊥_ (*η*=0), rotational velocity is non-zero. Finally, in our simulations, we have assumed that the grafting direction of two flagella at the basal ends, which are positioned exactly symmetrically with respect to the axis of symmetry of basal apparatus, is fixed over time. This angle also affects mean rotational velocity. All simulations in Figs. 7 and 8A-C (Videos 7-10) are done with V-angle of 180°, but we also performed simulations with V-angle of 90° which is closer to the experimentally measured values and observed reduction in mean rotational velocity (Figs. 8D-F and Videos 11-13).

**Figure 7.**
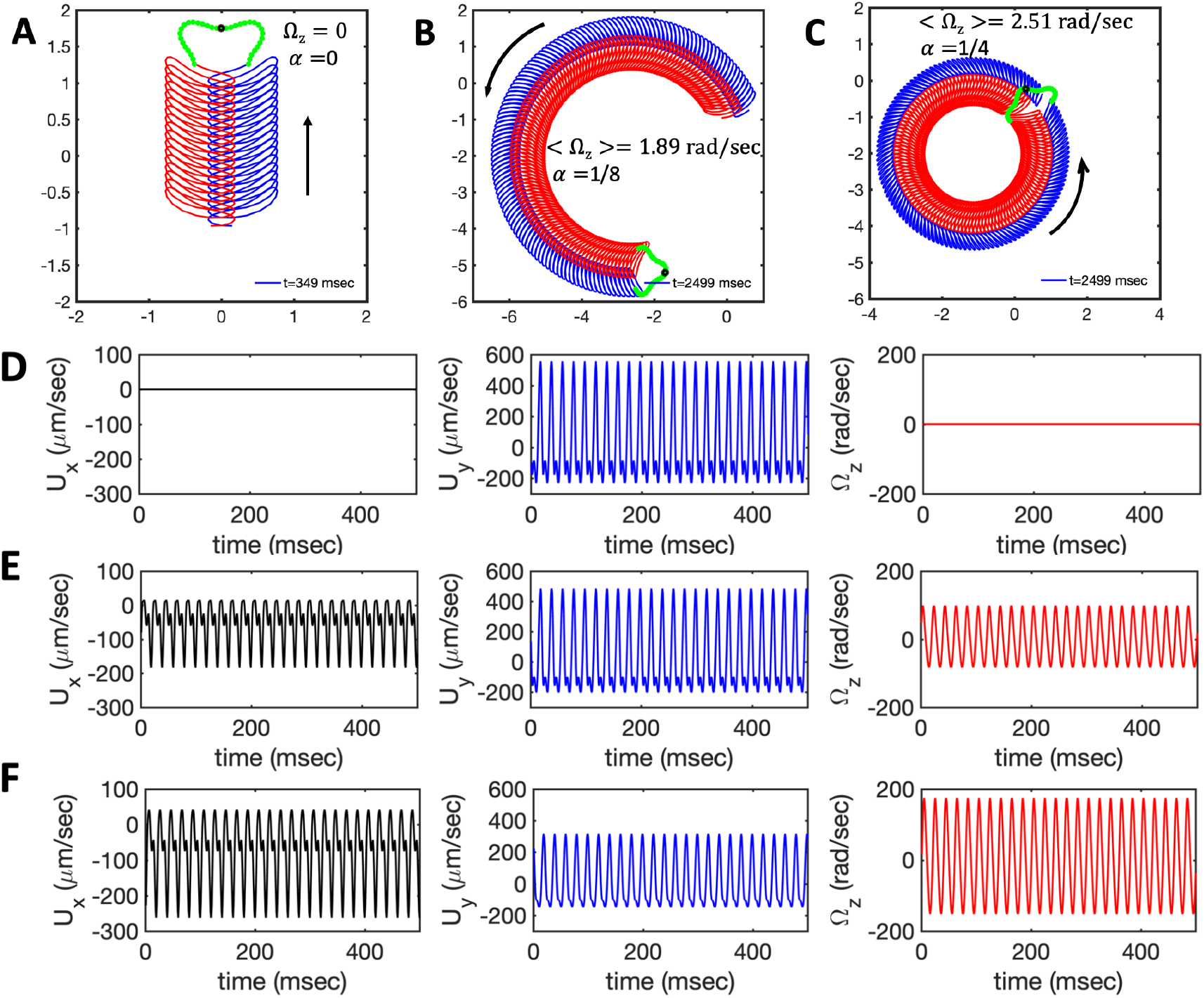
Simulations in the framework of RFT with a simplified wave pattern formed of superposition of a static and a dynamics mode. All parameters are the same for both flagella except the phase shift 2*πα*. A) Unsteady straight swimming for a flagellar apparatus with *a* = 0. The only non-zero component of velocity is *U_y_* which oscillates over time and there is no rotation. B) Circular swimming path for swimmer with *α* = 1/8 and C) *α* = 1/4. The mean rotational velocity is higher for larger phase shift. D-F) Instantaneous translational and rotational velocities in body-fixed frame corresponding to parts A, B and C, respectively. Note that both *U_x_, U_y_* and Ω_*z*_ oscillate at flagellar beat frequency of 50 Hz. Other parameters are: 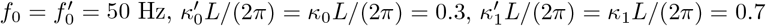. Note that parameters with prime are for flagella with blue trajectory (see Videos 5-7).

**Figure 8.**
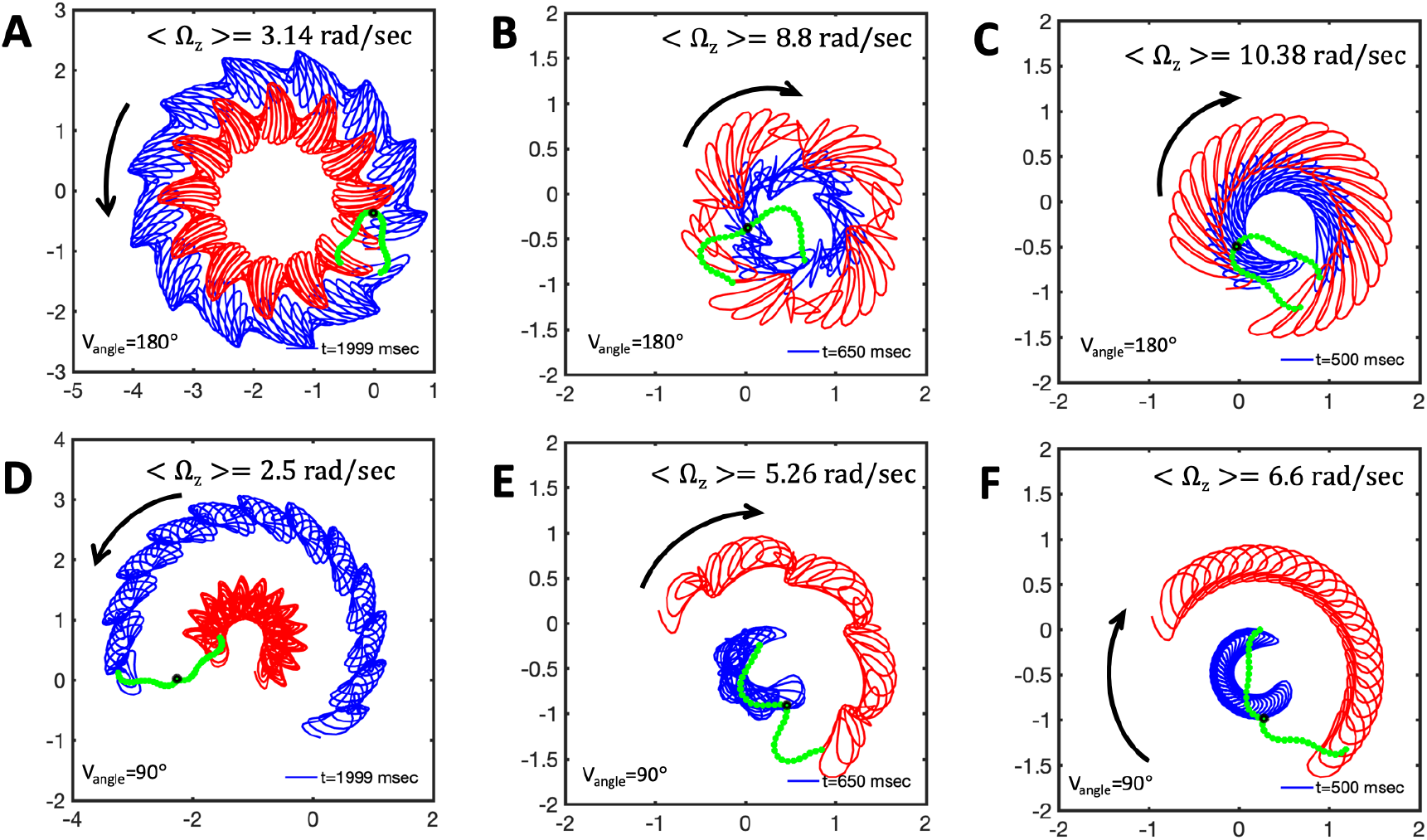
Simulations with simplified wave pattern to study the influence of variations in frequency and amplitude of the dynamic mode *κ*_1_. A) Flagella with blue trace beats faster compared to the flagellum with red trace (56 Hz versus 50 Hz) and therefore it sets the sign of (Ω_*z*_). B) To switch the direction of rotation, slower beating flagellum should beat with larger wave amplitude *κ*_1_. C) Keeping the frequencies and intrinsic curvature similar, flagellum with larger *κ*_1_ (red trace) sets the direction of swimming. Parameters are 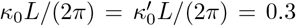, V-junction angle=180 °, *α* = 0 for all simulations, and A) 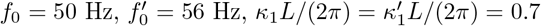, B) 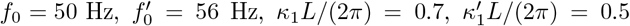, C) 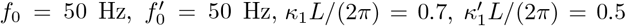. D-F) Simulations in parts A-C are repeated with V-junction angle of 90°. Parameters with prime are for flagella with blue trajectory (see Videos 8-13).

### 3.4. Discussion

In this paper, we have used high-speed imaging, quantitative image processing, and mode analysis to study wave dynamics of flagellar apparatus isolated from wall-less strain of *C.reinhardtii*. For isolation of this unique in vitro system, we followed the protocol established by Hyams & Borisy [32, 33], which has a very low yield; only 2-3% of the cells release their flagellar apparatus. In contrast to results reported in Refs. [32, 33], almost all of the isolated apparatus in our experiments had an intrinsic frequency mismatch around 4-6 Hz and therefore, it was not possible with our data to investigate synchronization dynamics in swimmers with no or very small frequency differences. We focused on freely swimming apparatus with intrinsic interflagellar frequency difference of Δ*f* ~ 4 Hz (~ 16% of the mean). In the presence of such a frequency mismatch, coupling between two flagella either via elastic fibers connecting the basal bodies, interflagellar hydrodynamic interactions or swimmer movement are not strong enough to overcome high frequency mismatch and the effect of noise (which acts against synchronization) to entrain two flagella. This is in agreement with experimental observations of intact *C. reinhardtii* cells [12] with high interflagellar frequency difference (10%-30%), supporting the hypothesis that intrinsic frequencies of flagella are controlled by signaling processes inside the cell.

We tracked waveforms of each flagellum using gradient vector flow technique [35, 36] with high spatio-temporal resolution, allowing us to characterize bending waves that propagate from basal ends towards the distal tips at frequency of 25±4 Hz. We performed mode analysis of flagella exhibiting rhythmic patterns [37] to define a continuous phase for each oscillator flagellum. Phase analysis captures the dynamics of two interacting flagella in basal apparatus, confirming that they effectively act as two isolated pendulums perturbing each other by non-linear fluctuations, but no entrainment can occur and interflagellar phase difference grows linearly with Δf (known as phase drift) [19, 24].

Further, we used our tracked data to examine the validity of resistive force theory for flagellar apparatus swimming in the vicinity of a planar substrate. From experimental recorded videos, we extracted the position of basal body with sub-micron resolution and calculated translational and rotational velocities. It is to be noted that as flagellar apparatus swims, angle of the V junction changes over time (see supplemental Videos 1-2). We computed instantaneous V-angle from tracked data and used it as an experimental input for our RFT analysis. Comparing our experimental data with instantaneous translational and rotational velocities of basal apparatus obtained by RFT simulations, we found a quantitative agreement with drag anisotropy of 2. Originally, the ratio ζ_⊥_/ζ_‖_ = 2 was used by Gray and Hancock for sea-urchin spermatozoa swimming far away from boundary [47]. In our experiments, flagellar apparatus swims in the vicinity of a glass surface and in this respect, our system is similar to experiments with microtubules moving parallel to a kinesin-coated substrate which can exert forces on microtubules [48]. Finally, we performed simulations and analytical approximations using simplified wave forms, allowing us to estimate mean rotational velocity of flagellar apparatus in the limit of small intrinsic curvature and wave amplitude. Interestingly, our analysis demonstrates that by introducing a phase or frequency difference between two flagella, steering of flagellar apparatus can occur.

It would be worthwhile to extend our analysis to flagellar apparatus without intrinsic frequency mismatch as experimentally observed in Refs. [32, 33]. Moreover, Hyams & Borisy report on interesting observation of switching swimming direction of basal apparatus from forward to backward motion at calcium concentrations above 1 *μ*M. In our experiments, with standard reactivation conditions, only forward swimming motion was observed. Calcium ions presumably affect the form and direction of ciliary beating patterns and it is known that exchange of calcium ions is crucial for tactic behavior of *C. reinhardtii* cells [33, 49]. Investigations in this direction are under way in our laboratory.

## Acknowledgments

We thank E. Bodenschatz, Y. Wang and A. Zaben for helpful comments and discussions and S. Romanowsky, M. Müller and K. Gunkel for technical assistance. A.G. and A.B. acknowledge MaxSynBio Consortium, which is jointly funded by the Federal Ministry of Education and Research of Germany and the Max Planck Society. The authors also thank M. Lorenz and the Göttingen Algae Culture Collection (SAG) for providing the *Chlamydomonas reinhardtii* wall-less strain SAG 83.81.

## Supplemental Information

## Appendix I: Approximation of mean rotational velocity

First, we consider the dynamics of a single flagellum. In the simplified form, curvature waves propagating along the contour length of a flagellum is given by:

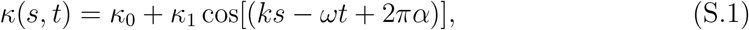

where *ω* = 2*π f*_0_ and *k* = 2*π*/λ ~ 2*π/L*. Tangential angle *θ* is computed as:

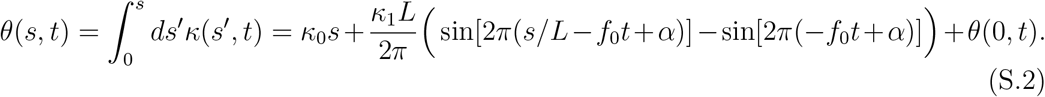

For simplicity, we assume flagella is clamped at *s* = 0 along the 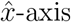, i.e. *θ*(0, t) = 0 and *x*(*s* = 0, *t*) = *y*(*s* = 0, *t*) = 0. Assuming both *κ*_0_/*k* ~ *κ*_0_*L*/(2*π*) and *κ*_1_/*k* ~ *κ*_1_*L*/(2*π*) to be small, we approximate cos *θ* and sin *θ* by keeping *κ*_0_*L*/(2*π*) up to the second order and *κ*_1_*L*/(2*π*) up to the third order:

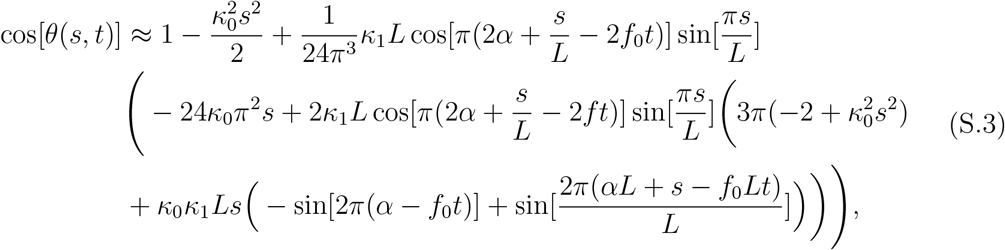

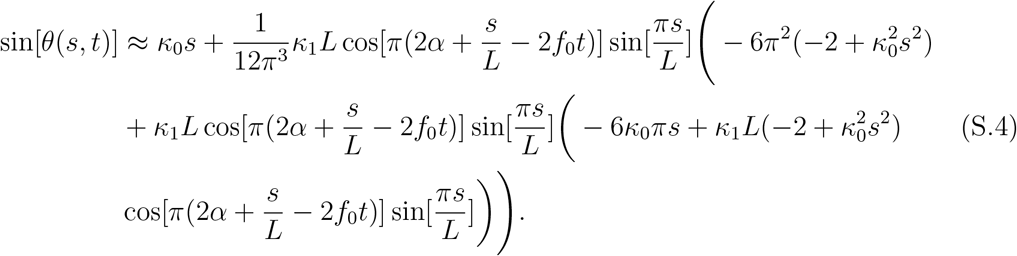

Flagella’s centerline is described by vector 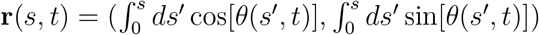 and local velocity and unit tangent vectors are calculated as **U**(*s,t*) = (*dx/dt, dy/dt*) and **t** = (**t**_*x*_, **t**_*y*_) = (cos[*θ*(*s, t*)], sin[*θ*(*s, t*)]), respectively.

We calculate instantaneous active torque around the grafting point at (0, 0) as:

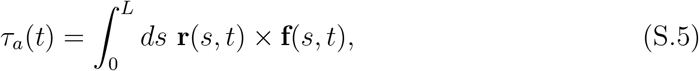

where we **f**(*s, t*) is computed in the framework of resistive force theory:

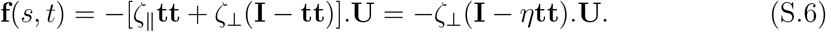

Here **tt + nn = I** is the identity matrix and *η* = (ζ_⊥_ − ζ_‖_)/*ζ*_⊥_ is a measure of anisotropy in drag coefficients. Note that if we calculate time-average of force **f**(*s, t*), the first term in Eq. S.6 vanishes and only the second term which is proportional to drag anisotropy contributes in the mean force. However, the first term does not vanish when we calculate the mean torque. Thus, *τ_a_*(*t*) has a term proportional to ζ_⊥_ and a second term proportional to ζ_⊥_*η*:

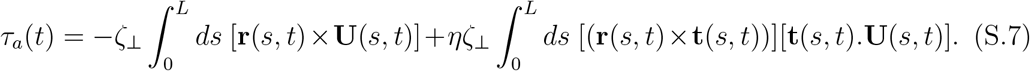

For a planar flagellar beat in *x − y* plane, *τ_α_*(*t*) is in *z* direction. In the limit of λ → *L* and up to second order in *κ*_0_*L*/(2*π*) and third order in *κ*_1_*L*/(2*π*), we obtain:

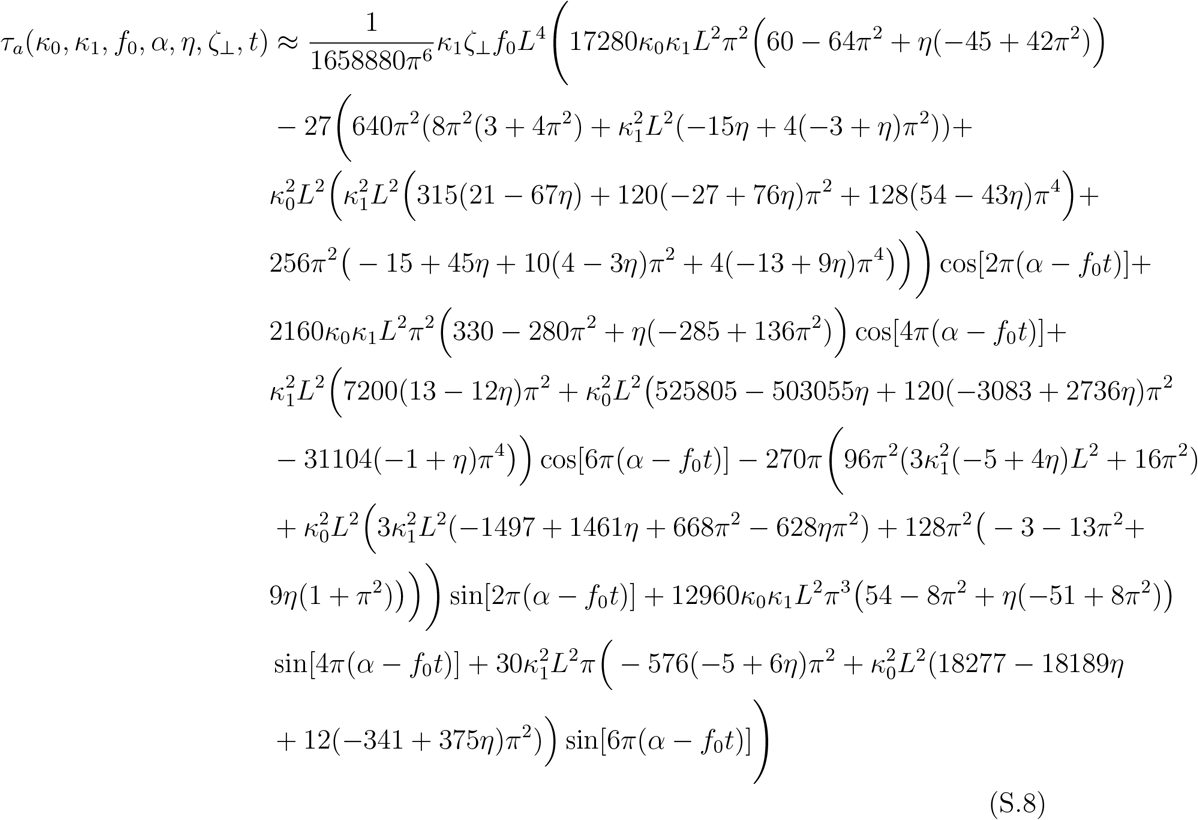

As mentioned previously, even in the absence of drag anisotropy (*η* = 1 − ζ_‖_/ζ_⊥_ = 0), Eq. S.8 has a non-zero value proportional to ζ_⊥_.

In parallel, we approximate the instantaneous viscous torque exerted on a flagellum which deforms its shape over time while it is pinned at one end and rotates at instantaneous angular velocity Ω = (0, 0, Ω_*z*_(*κ*_0_, *κ*_1_, *f*_0_, *α, η, t*)) around the pinning point. At any instant of time, we consider flagella apparatus as a solid body with rotational velocity Ω_*z*_ and calculate the viscous torque as:

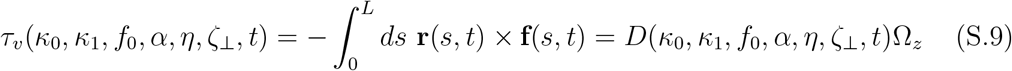

where **f**(*s, t*) is calculated using Eq. S.6 with instantaneous rigid body velocity 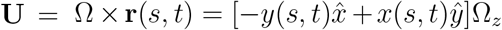 and *D* is the instantaneous drag coefficient that we aim to calculate. After performing the integration in Eq. S.9, we obtain instantaneous drag coefficient as:

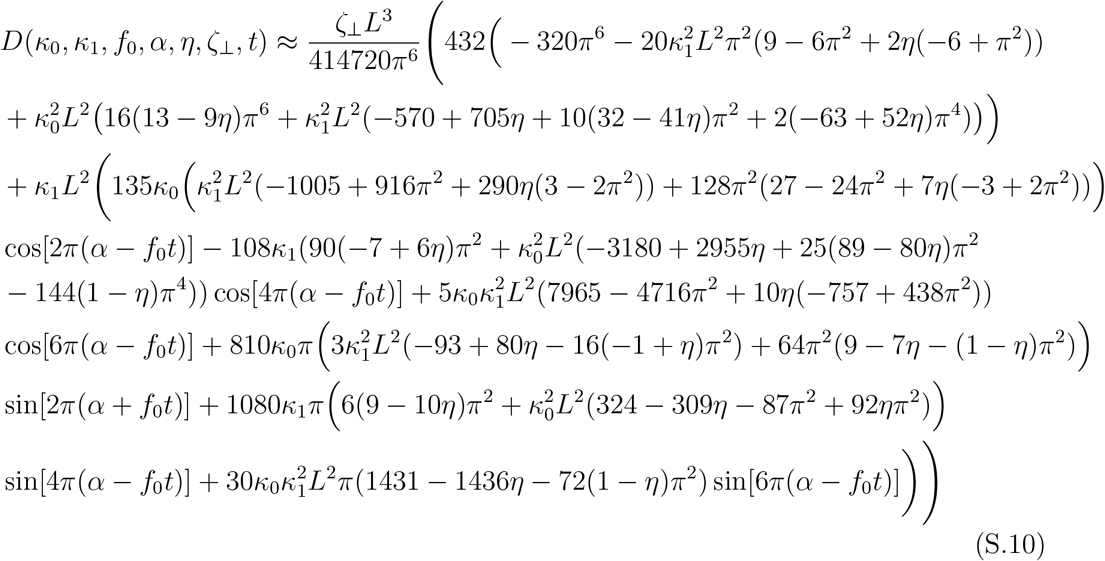

Note that for *κ*_0_ = *κ*_1_ = 0, Eq. S.10 reduces to 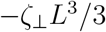 which is the drag coefficient of a rigid cylinder of length *L*. Furthermore, in the limit of κ_1_ = 0, Eq. S.10 simplifies to 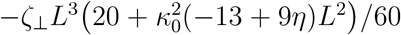 which is the drag coefficient of a bent cylinder with mean curvature *κ*_0_.

Now, we can estimate mean rotational velocity for a single flagellum as:

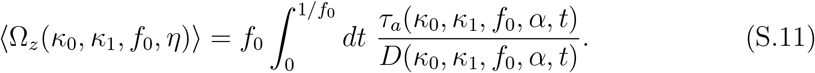

Before performing the integration over time, we expand the ratio of *τ_a_/D* up to first order in *κ*_0_*L*/(2*π*) and second order in *κ*_1_*L*/(2*π*), to obtain:

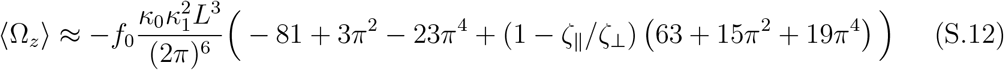

As expected, by integrating over one beat cycle, *α* averages out. Moreover, consistent with our simulations, (Ω_*z*_) is non-zero for isotropic drag coefficients ζ_‖_ = ζ_⊥_.

After estimating *τ_α_, τ_υ_* and (Ω_*z*_) for a single flagellum, we can now calculate instantaneous rotational velocity Ω_*z*_ of a flagellar apparatus with two flagella, ignoring the viscous drag of basal body for simplicity. We balance instantaneous total active torque with instantaneous total drag torque exerted on flagella apparatus to obtain:

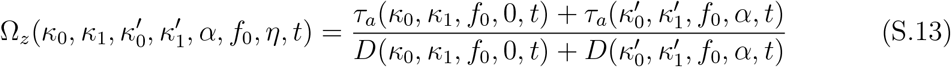

Note that two flagella have different values of intrinsic and dynamic curvature and beat with a phase difference of 2*πα*. Next, to estimate time-averaged rotational velocity (Ω_*z*_) of flagellar apparatus, we integrate over one beating cycle:

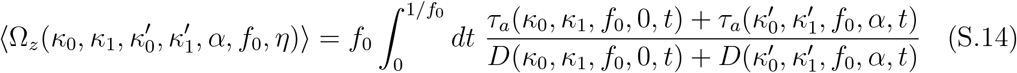

Here, for simplicity we have assumed that both flagella beat at the same frequency *f*_0_. Before doing the integration, we expand the ratio of total active torque to instantaneous drag coefficients up to first order in *κ*_0_*L*/(2*π*) and 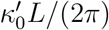 but second order in *κ*_1_*L*/(2*π*) and 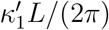 to obtain:

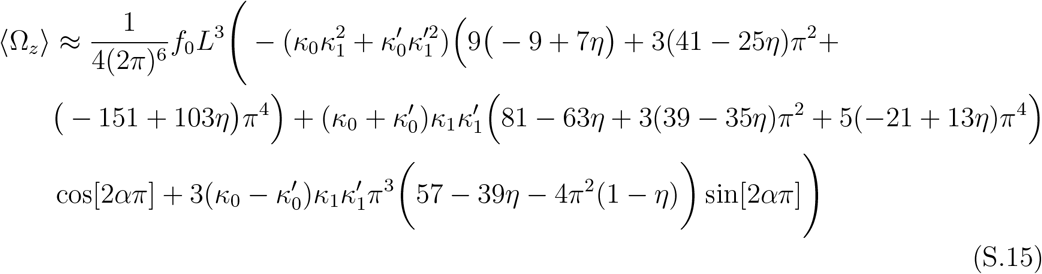

Remarkably, if both flagella have the same magnitude of intrinsic curvature 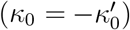 and beat at equal amplitude of dynamic mode 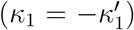 with phase difference 2*πα*, then Eq. S.15 will be reduced to Eq. 17 and in the limit of *α* = 0, it simplifies to Eq. 18.

## Supplemental Movies

**Video 1** Tracked trajectories plotted on top of experimental data for the first 800 msec.

**Video 2** Trajectory of basal body obtained by a Gaussian fit showing a helical path.

**Video 3** Video showing superposition of four eigenmodes on top of tracked flagellum.

**Video 4** Swimming trajectory of basal apparatus obtained by RFT simulations.

**Video 5** Simulations with simplified wave form with equal values of dynamic and static modes and frequency for both flagella and no phase difference (see Fig. 7A).

**Video 6** Simulations with simplified wave form with equal values of dynamic and static modes and frequency but phase difference of *π*/4 (see Fig. 7B).

**Video 7** Simulations with simplified wave form with equal values of dynamic and static mode but phase difference of *π*/2 (see Fig. 7C).

**Video 8** Simulations with simplified wave form with equal phase, *κ*_0_, *κ*_1_, but difference in frequency (see Fig. 8A).

**Video 9** Simulations with simplified wave form with equal phase and *κ*_0_, but different frequency and *κ*_1_ (see Fig. 8B).

**Video 10** Simulations with simplified wave form with equal phase, frequency and *κ*_0_, but different *κ*_1_ (see Fig. 8C).

**Video 11** Same as Video 8 with V_angle_ = 90° (see Fig. 8D).

**Video 12** Same as Video 9 with V_angle_ = 90° (see Fig. 8E).

**Video 13** Same as Video 10 with V_angle_ = 90° (see Fig. 8F).

## Supplemental Figures

**Figure S. 1.**
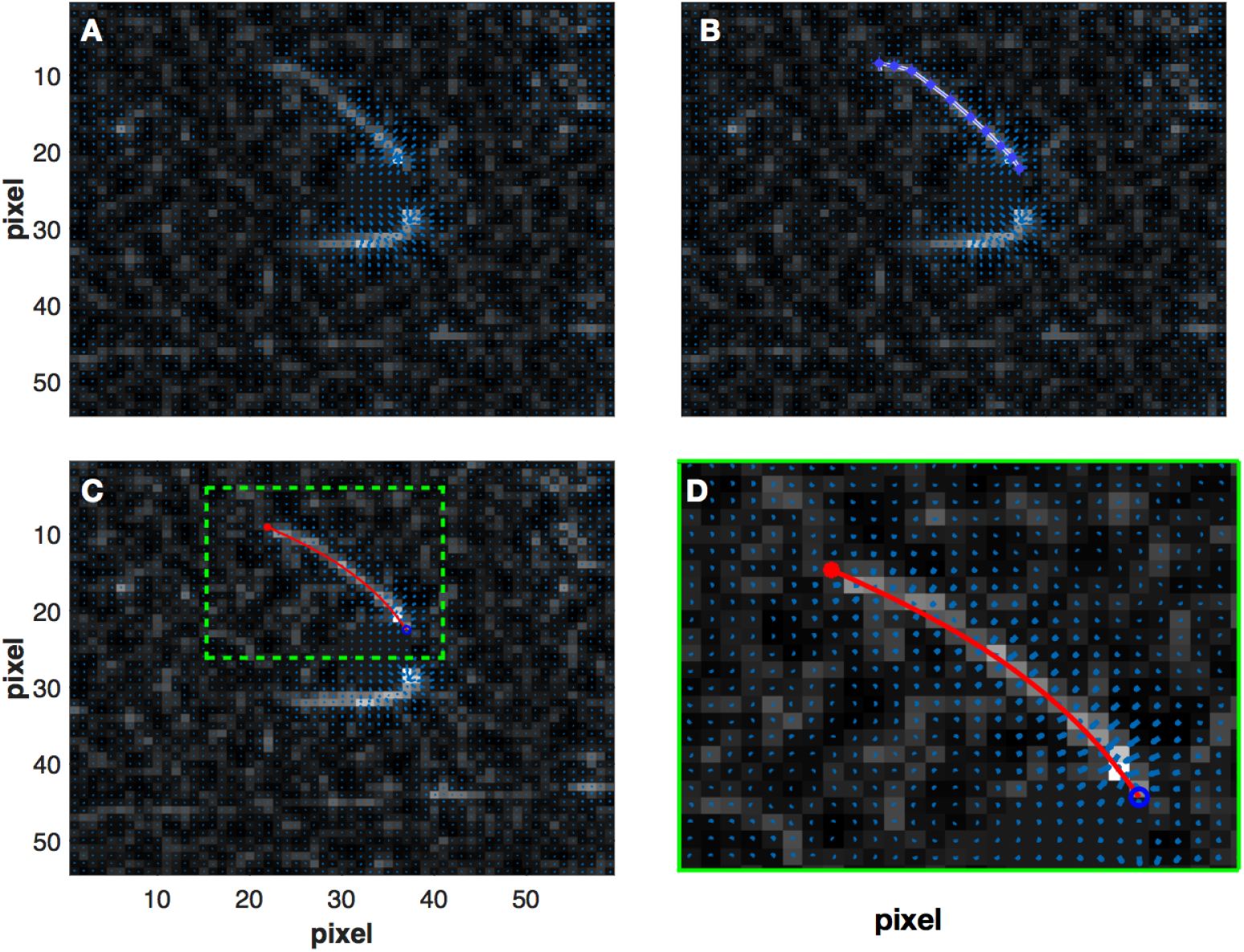
A) Gradient vector flow (blue arrows) calculated around flagella apparatus. B) The initial selection of a polygon for the first frame which deforms according to the gradient vector field calculated around the top flagellum. C) The final tracked shape of top flagella. D) A zoomed-in image showing GVF in higher magnification in the vicinity of top flagella.

## Reference

[1] Eggenschwiler J T and Anderson K V 2007 Annu. Rev. Cell Dev. Biol. 23 345–373

[2] Marshall W F and Nonaka S 2006 Current Biology 16 R604–R614

[3] Goetz J G, Steed E, Ferreira R R, Roth S, Ramspacher C, Boselli F, Charvin G, Liebling M, Wyart C, Schwab Y et al. 2014 Cell reports 6 799–808

[4] Machemer H 1972 Journal of Experimental Biology 57 239–259

[5] Goldstein R E 2015 Annual review of fluid mechanics 47 343–375

[6] Gilpin W, Bull M S and Prakash M 2020 Nature Reviews Physics 1–15

[7] Faubel R, Westendorf C, Bodenschatz E and Eichele G 2016 Science 353 176–178

[8] Kramer-Zucker A G, Olale F, Haycraft C J, Yoder B K, Schier A F and Drummond I A 2005 Development 132 1907–1921

[9] Mitchison T and Mitchison H 2010 Nature 463 308–309

[10] Burgess S A, Walker M L, Sakakibara H, Knight P J and Oiwa K 2003 Nature 421 715–718

[11] Ishikawa T, Sakakibara H and Oiwa K 2007 Journal of molecular biology 368 1249–1258

[12] Polin M, Tuval I, Drescher K, Gollub J P and Goldstein R E 2009 Science 325 487–490

[13] Goldstein R E, Polin M and Tuval I 2009 Physical review letters 103 168103

[14] Taylor G I 1951 Proceedings of the Royal Society of London. Series A. Mathematical and Physical Sciences 209 447–461

[15] Uchida N and Golestanian R 2011 Physical Review Letters 106 058104

[16] Golestanian R, Yeomans J M and Uchida N 2011 Soft Matter 7 3074–3082

[17] Vilfan A and Jülicher F 2006 Physical review letters 96 058102

[18] Lauga E and Powers T R 2009 Reports on Progress in Physics 72 096601

[19] Friedrich B 2016 The European Physical Journal S’pecial Topics 225 2353–2368

[20] Bennett R R and Golestanian R 2013 Physical review letters 110 148102

[21] Quaranta G, Aubin-Tam M E and Tam D 2015 Physical review letters 115 238101

[22] Geyer V F, Jülicher F, Howard J and Friedrich B M 2013 Proceedings of the National Academy of Sciences 110 18058–18063

[23] Friedrich B M and Jülicher F 2012 Physical Review Letters 109 138102

[24] Wan K Y, Leptos K C and Goldstein R E 2014 Journal of the Royal Society Interface 11 20131160

[25] Ringo D L 1967 The Journal of cell biology 33 543–571

[26] Lechtreck K F and Melkonian M 1991 An update on fibrous flagellar roots in green algae The Cytoskeleton of Flagellate and Ciliate Protists (Springer) pp 38–44

[27] Wan K Y and Goldstein R E 2016 Proceedings of the National Academy of Sciences 113 E2784–E2793

[28] Harris E H 2001 Annual review of plant biology 52 363–406

[29] Rüffer U and Nultsch W 1985 Cell Motility 5 251–263

[30] Sleigh M 1979 Nature 277 263–264

[31] Klindt G S, Ruloff C, Wagner C and Friedrich B M 2017 New Journal of Physics 19 113052

[32] Hyams J S and Borisy G G 1975 Science 189 891–893

[33] Hyams J S and Borisy G G 1978 Journal of Cell Science 33 235–253

[34] Geyer V 2013

[35] Xu C and Prince J L 1997 Gradient vector flow: A new external force for snakes *IEEE Proc*. CVPR (IEEE) pp 66–71 ISBN 0-8186-7822-4

[36] Xu C, Prince J L et al. 1998 IEEE Trans. Image Proc. 7 359–369

[37] Stephens G J, Johnson-Kerner B, Bialek W and Ryu W S 2008 PLoS computational biology 4

[38] Keller J B and Rubinow S 1976 Biophysical Journal 16 151–170

[39] Johnson R and Brokaw C 1979 Biophysical journal 25 113–127

[40] Leach J, Mushfique H, Keen S, Di Leonardo R, Ruocco G, Cooper J and Padgett M 2009 Physical Review E 79 026301

[41] Saggiorato G, Alvarez L, Jikeli J F, Kaupp U B, Gompper G and Elgeti J 2017 Nature communications 8 1–9

[42] Sartori P, Geyer V F, Scholich A, Jülicher F and Howard J 2016 Elife 5 e13258

[43] Geyer V F, Sartori P, Friedrich B M, Jülicher F and Howard J 2016 Current Biology 26 1098–1103

[44] Friedrich B M, Riedel-Kruse I H, Howard J and Jülicher F 2010 Journal of Experimental Biology 213 1226–1234

[45] Lauga E 2007 Physical Review E 75 041916

[46] Shapere A and Wilczek F 1987 Physical Review Letters 58 2051

[47] Gray J and Hancock G 1955 Journal of Experimental Biology 32 802–814

[48] Hunt A J, Gittes F and Howard J 1994 Biophysical journal 67 766

[49] Bessen M, Fay R B and Witman G B 1980 The Journal of Cell Biology 86 446–455

